# Tonic and phasic transcutaneous auricular vagus nerve stimulation both evoke rapid and transient pupil dilation

**DOI:** 10.1101/2023.09.20.558607

**Authors:** Lina Skora, Anna Marzecová, Gerhard Jocham

## Abstract

**Background:** Transcutaneous auricular vagus nerve stimulation (tVNS or taVNS) is a non-invasive method of electrical stimulation of the afferent pathway of the vagus nerve, suggested to drive changes in putative physiological markers of noradrenergic activity, including pupil dilation.

**Objective:** However, it is unknown whether different taVNS modes can map onto the phasic and tonic modes of noradrenergic activity. The effects of taVNS on pupil dilation in humans are inconsistent, largely due to differences in stimulation protocols. Here, we attempted to address these issues.

**Methods:** We investigated pupil dilation under phasic (1 s) and tonic (30 s) taVNS, in a pre-registered, single-blind, sham-controlled, within-subject cross-over design, in the absence of a behavioural task.

**Results:** Phasic taVNS induced a rapid increase in pupil size over baseline, significantly greater than under sham stimulation, which rapidly declined after stimulation offset. Tonic taVNS induced a similarly rapid (and larger than sham) increase in pupil size over baseline, returning to baseline within 5 s, despite the ongoing stimulation. Thus, both active and sham tonic modes closely resembled the phasic effect. There were no differences in tonic baseline pupil size, and no sustained effects of stimulation on tonic baseline pupil size.

**Conclusions:** These results suggest that both phasic- and tonic-like taVNS under the standard stimulation parameters may modulate primarily the phasic mode of noradrenergic activity, as indexed by evoked pupil dilation, over and above somatosensory effects. This result sheds light on the temporal profile of phasic and tonic stimulation, with implications for their applicability in further research.

## Introduction

Transcutaneous auricular vagus nerve stimulation (tVNS or taVNS) is a non-invasive method of electrical stimulation of the vagal afferent pathway delivered transcutaneously to the auricular branch of the vagus nerve (ABVN) [1–4] by placing the electrodes on the skin of the outer ear innervated by the ABVN. Recent evidence suggests that taVNS can serve as a tool for modulating noradrenaline (NE) levels [5]. In addition to promising results in treatment of pharmaco-resistant depression and epilepsy [6–8], taVNS has been proposed as a non-invasive tool to probe neuromodulatory influences on human behaviour [9,10]. However, the effect of taVNS on markers of noradrenergic activity is still unclear, with different studies yielding an inconsistent pattern of results. This is likely a consequence of the variety of combinations of stimulation parameters that have been used (particularly intensity, pulse width, frequency, duration of stimulation) [9,11]. Given the limited understanding of the precise mechanism of action, a clearer mapping of taVNS-induced changes in potential markers of noradrenergic activity is imperative.

The afferent vagus nerve has highly diffuse targets in the brain, complicating the understanding of the neuromodulatory mechanisms of taVNS. The nerve terminates at the nucleus of the solitary tract (NTS), projecting to the parabrachial nucleus, locus coeruleus (LC; the primary source of NE in the brain), as well as the dorsal raphe nucleus (the primary source of serotonin) in the brainstem, and further to the periaqueductal grey, thalamus, nucleus accumbens, amygdala, insula, and hippocampus [3,12,13]. Those targets have been found to be activated by taVNS in sham-controlled human neuroimaging studies [3,4,14–17].

Stimulation with implanted VNS (iVNS) in rodents has been shown to robustly elicit LC firing and increase noradrenergic discharge, in a dose-dependent manner [5,18,19]. Two modes of LC-NE firing patterns have been described: the tonic mode, characterised by a continuous, low-frequency firing pattern, and phasic mode, characterised by short bursts at higher frequencies [20]. The phasic mode, driven by task-related stimuli, has been proposed to optimise task engagement, while the tonic mode has been linked to attentional disengagement and exploration [20]. It is currently unclear whether and to what extent taVNS modes and parameters can map onto the two modes of LC-NE discharge. In addition, the picture is obscured by inconsistencies in stimulation protocols and the resulting biomarkers (see review by [9]).

In the current study, we focus on pupil diameter as a marker of LC-NE activity. Pupil dilation is modulated by sympathetic and parasympathetic influences [21], and, when not mediated by light levels, is suggested to be driven primarily through activation of the LC, thought to increase activity in the dilator muscle while inhibiting activity in the iris sphincter [22–24]. Using pupil size as a putative marker of noradrenergic activity in humans [25,26], taVNS applied at the standard clinical duty protocol of 30 s on/30 s off typically results in an absence of a tonic change in pupil size [27–32] (see Table 1), although contrary results have been reported both for taVNS [33] and human iVNS [34]. In contrast, in more recent studies with short, phasic-like (500ms - 2 min) stimulation pulses, taVNS has been found to elicit a rapid pupil dilation, peaking and returning to baseline within a few seconds from stimulation onset [11,35–38]. A comparable effect has been found in rodents with iVNS [18,39,40]. Together, these results suggest that stimulating the LC through the afferent vagal projections might induce a phasic noradrenergic discharge reflected in rapid and short-lived pupil dilation. Nonetheless, the temporal profiles of tonic and phasic-like taVNS under matched parameters have not yet been compared.

**Table 1.**
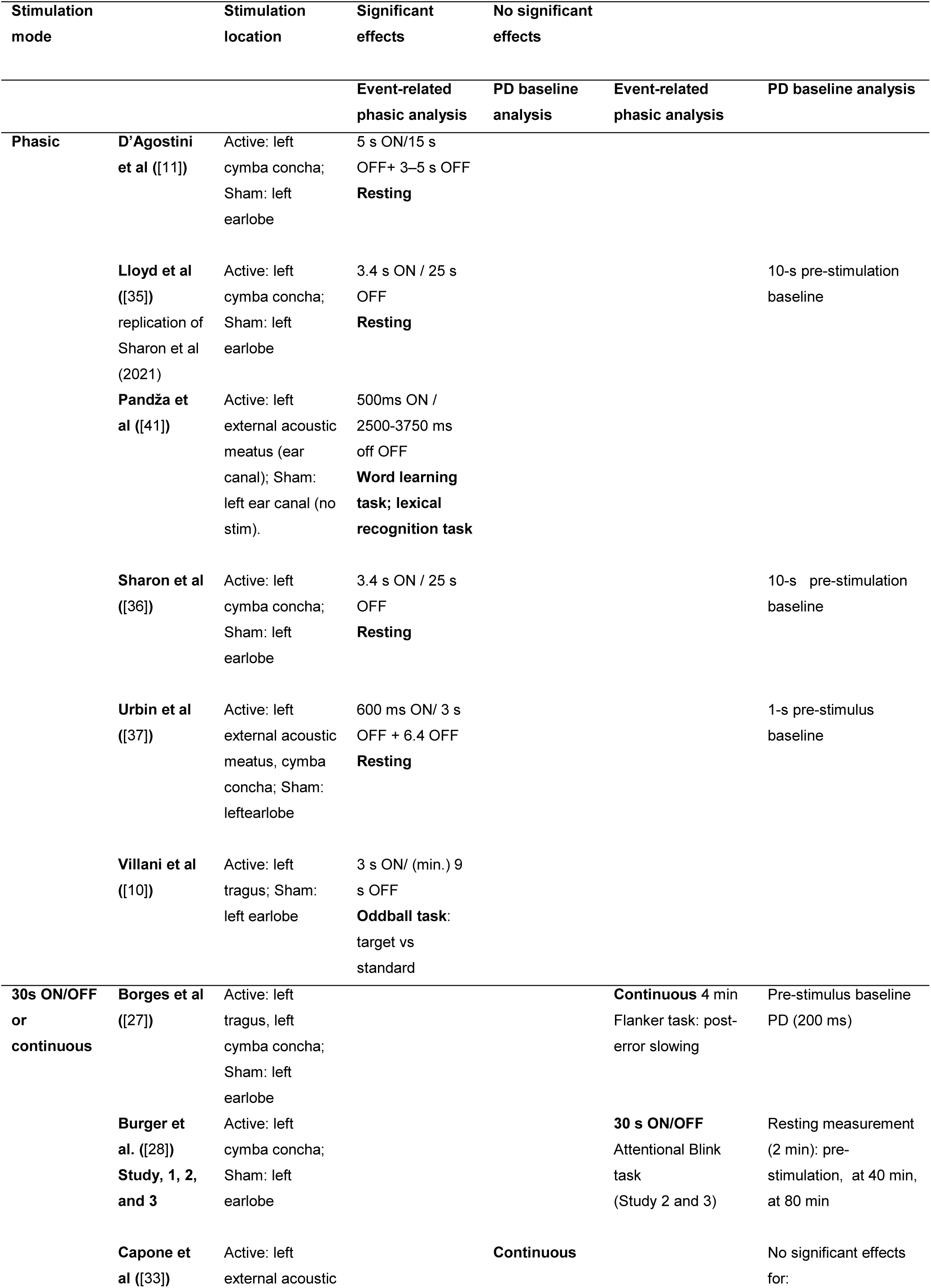
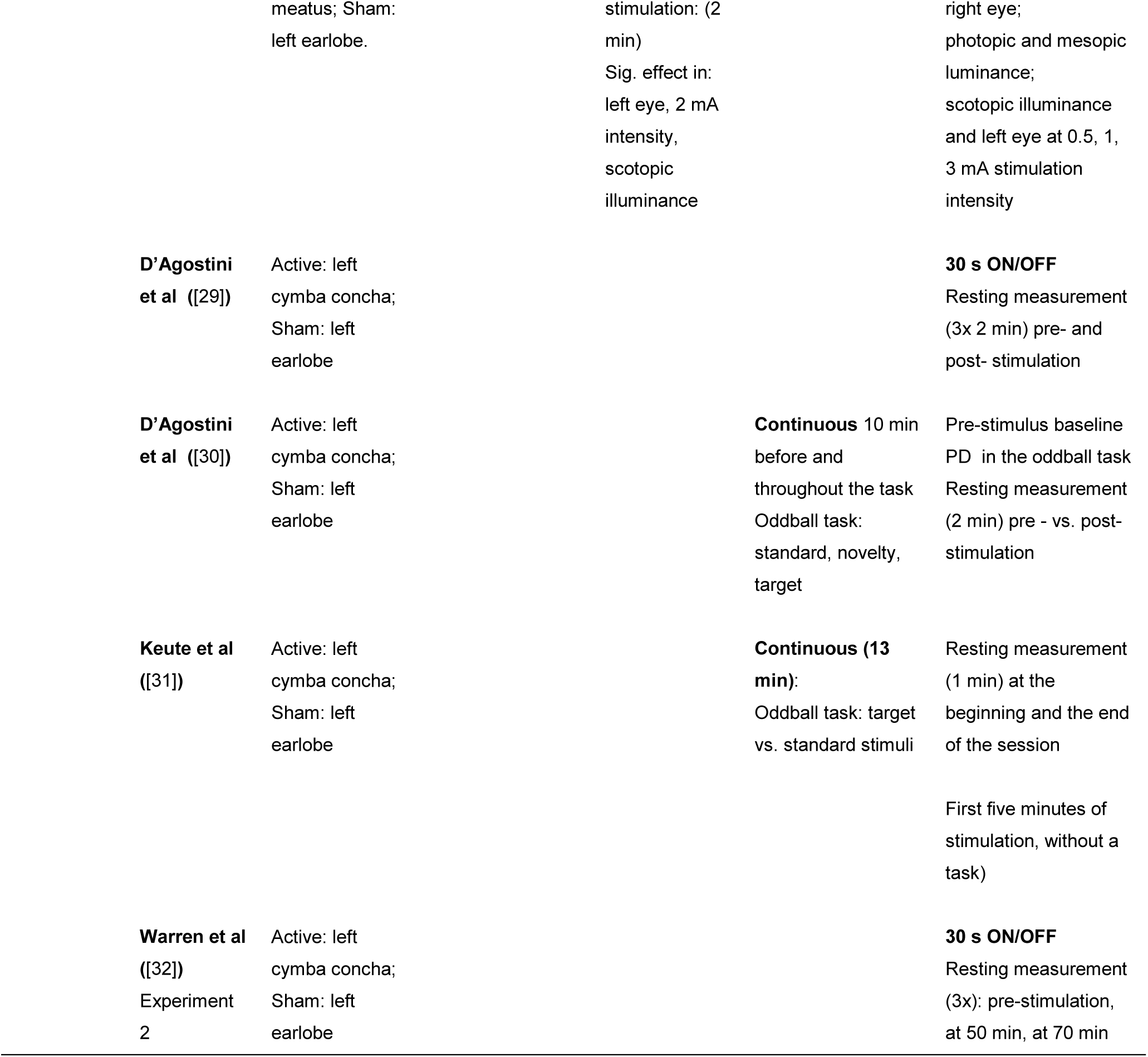
Overview of past evidence on the effects of taVNS on pupil diameter (PD) across studies investigating phasic-like (brief, 0.6-5 s) stimulation and prolonged, tonic-like (30s ON/OFF or continuous) stimulation.

Here, we set out to investigate pupil dilation under phasic (1 s) and tonic (30 s) stimulation, in a single-blind, sham-controlled, within-subject cross-over design, in the absence of a behavioural task. To our knowledge, the present study is the first to compare phasic and tonic stimulation protocols with matched parameters in a within-subject design at rest, thus elucidating the temporal profiles of pupil diameter changes under both stimulation modes. Our secondary motivation was to use pulses as short as technically possible in the phasic condition, in order to explore their efficacy as a phasic noradrenergic manipulation, which could be used in behavioural tasks. We expected to replicate the previous findings that short-burst taVNS evokes a phasic event-related response. Furthermore, if the tonic stimulation protocol elevates tonic LC activity reflected in baseline pupil values, we expected to observe larger tonic baseline pupil values for active vs sham stimulation. Finally, if tonic stimulation, similarly to phasic stimulation, elicits a transient event-related response, we hypothesised the temporal profile of pupil dilation under tonic stimulation to resemble that under phasic stimulation. This approach can shed light on the efficiency of taVNS as a noradrenergic modulatory tool, as well as its potential for use in an event-related manner for behavioural tasks.

## Method

### Participants

61 participants (38 females, 2 other/prefer not to say; *M*_AGE_ = 22.3, range = 18-34, age was missing for 4) were recruited to participate among the student population of Heinrich Heine University Düsseldorf. Participants were native German speakers, or spoke fluent or native English. Sample size of 52 was determined to be necessary for obtaining a small-to-medium effect size with 80% power (computed with G*Power [42] for the Active v Sham comparison), but extra participants were collected in order to account for possible drop-outs or exclusions. Three participants failed to complete one or both sessions, yielding 58 analysable participants. Participants were compensated at the rate of 10 EUR/h, with a 10 EUR bonus for attending both testing sessions (50 EUR total).

All participants had normal or corrected-to-normal vision. Participants reported no current or history of neurological or psychiatric disease, cardiovascular disease, arrhythmia, seizures, hypertension, head injury, hearing problems, vision problems, mobility problems, thyroid dysfunction, diabetes, asthma, epilepsy, glaucoma, haemophilia, impaired liver or kidney function, and no substance abuse. They had no cochlear implants, cardiac pacemakers, implanted vagus nerve stimulators (VNS), or cerebral shunts, and reported no current sores, dryness, or skin disorders (e.g. eczema) on the ears. They were required to take no prescription medication, centrally-acting or known to increase seizure risk (e.g. buproprion, neuroleptics, albuterol, theophylline, antidepressants, thyroid medication, or stimulants). The study was approved by the Ethics Committee of the Medical Department at the HHU (ref. 2021-1754_1).

The study was pre-registered on OSF at https://osf.io/6gsfp.

### Stimuli and Equipment

The experiment was conducted in Matlab (Mathworks, 2019b), running Psychtoolbox [43]. The task was presented on a 24 inch Asus PG248Q monitor (1440 by 900 resolution). The only stimulus used was a single black-and-white fixation point combining a bullseye and cross-hair (0.6 x 0.2 degrees of visual angle), generated following Thaler et al [44] recommendation to ensure stable central fixation and minimise eye movements. The fixation point was presented on a grey screen (RGB 100,100,100). The task code can be found at https://osf.io/v2u9s/.

#### Eyetracking

Throughout the task, participant’s pupil size, as well as horizontal and vertical coordinates of eye position from both eyes were recorded with an Eyelink1000 Plus [45] eye tracker in arbitrary units (a.u.). For a portion of participants, the data were recorded as diameter, while for another portion, data were recorded as area, and subsequently converted (see *Data processing*). Data were sampled at 500 Hz (note that in 25% of participants data were sampled at a higher rate and downsampled prior to pre-processing), at 75% (in the majority of participants) or 100% illuminator power. Participants’ heads were maintained in a stable position using a chin and forehead rest placed 78 cm from the screen, in a room with constant, dim, ambient light. No images of the pupils were collected.

*taVNS:* A tVNS® R [46] taVNS device with legacy electrodes was used to stimulate the left auricular branch of the vagus nerve (ABVN). Two titanium electrodes, covered in a conductive pad and coated in conductive cream, were placed in the cymba conchae of the left ear for active stimulation, or on the left earlobe for sham stimulation. Particularly in the sham position, where electrode-skin contact was more challenging to maintain, electrodes were kept in place with medical tape and a headband (note that the headband was used for some participants for whom the electrode would have been attached less securely than in the active condition, see Fig.1). Electrical stimulation was delivered as a biphasic square wave, with impulse frequency of 25 Hz and pulse width of 250 μs.

**Figure 1.**
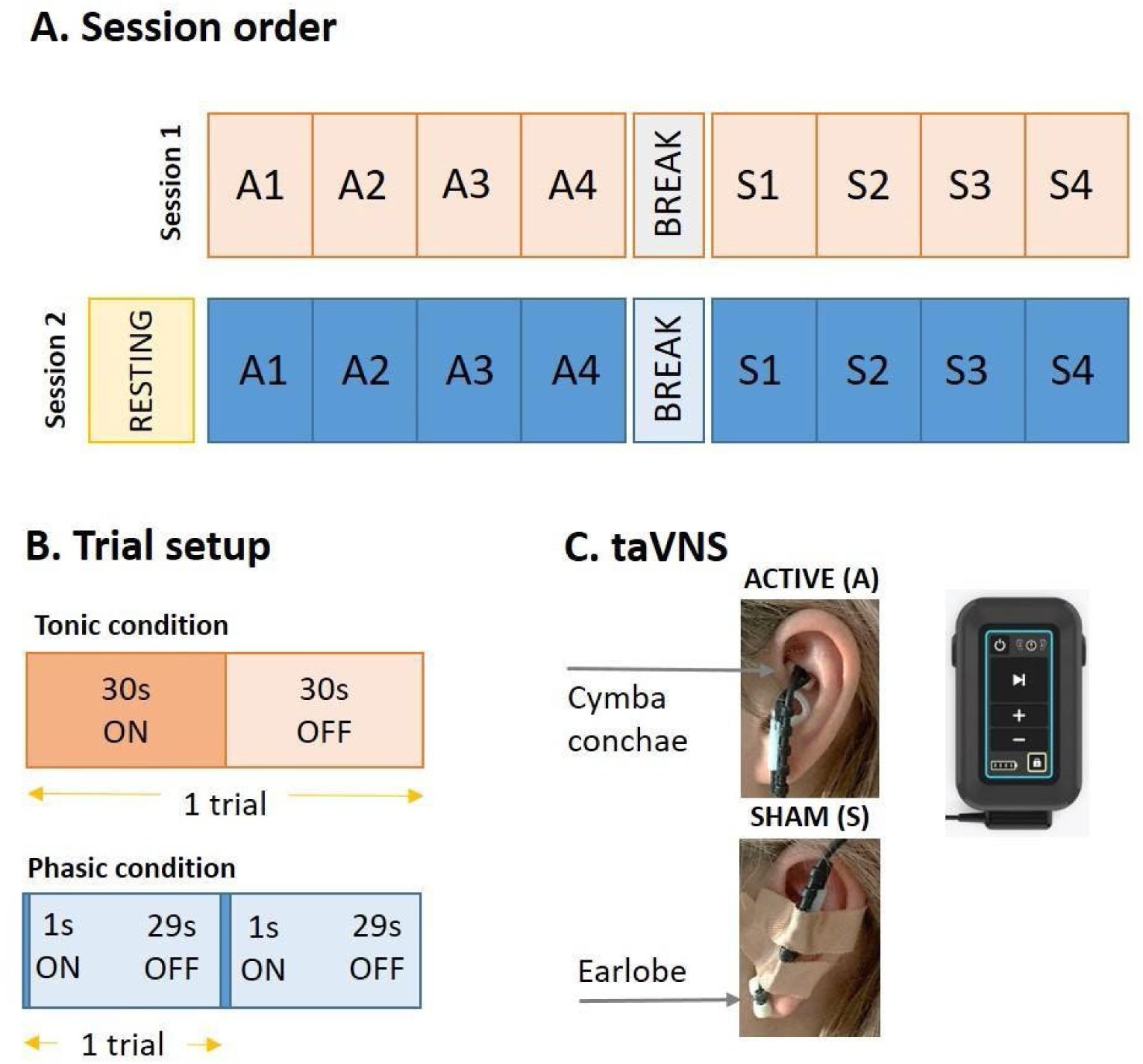
**A:** Example of session order and stimulation mode order in the sessions. Both the session order (e.g. phasic in session 1, tonic in session 2) and stimulation mode order (Active (A) first or Sham (S) first) were counterbalanced across subjects. **B:** Trial set-up. In the tonic condition, stimulation was delivered for 30 s (in both A and S modes), followed by 30 s off-stimulation, resulting in six 60 s long trials in each of the four blocks. In the phasic condition, stimulation was delivered for 1 s (in both A and S modes), followed by 29s off-stimulation, resulting in 11 30 s long trials in each of the four blocks. **C:** Stimulation locations on the left ear for Active (electrodes placed in the cymba conchae) and Sham (electrodes placed on the earlobe and secured with medical tape) modes (photo credit: authors). taVNS® R device (photo reproduced with permission by tVNS Technologies, Erlangen, Germany).

Stimulation intensity was calibrated individually for every participant with a work-up procedure to find a clearly perceptible, but not painful, sensation, within the range of 1-5 mA. Calibration was performed in the active mode (left cymba concha) at the beginning of the first session. In the calibration procedure, stimulation was manually started at the intensity of 1 mA, and increased in steps of 0.2 mA. After each stepwise increase, participants were asked to report their experience on a visual analogue scale from 1 (“I don’t feel it at all/Ich fuehle nichts”) to 5 (“schmerzhaft”/“painful”). The process continued until participants reached the rating of 5, in which case intensity was decreased to a level rated as 4 (“perceptible but not painful”). If the decreased intensity was rated as lower than 4, the stepwise increase again continued to a level rated as 5, in which case it was reduced until participants reported a sensation corresponding to level 4.

During the task, stimulation commands were sent to the tVNS® R device using a custom tVNS Technologies app (tVNS Manager, version 0.9.3) via a Bluetooth connection. The timings of stimulation onsets and offsets were controlled automatically from the Psychtoolbox/Matlab task by sending http POST requests to tVNS Manager, and were recorded in the Eyelink data file. Stimulation duration depended on the phasic or tonic condition (see *Procedure),* and was applied with immediate full-intensity onset (without ramp-up), which was critical for maintaining the 1 s stimulation duration. It also allowed for precise time-stamping of stimulation onsets. Onset of each trial was time-stamped after the completion of the start request. End of each trial was timestamped after the defined off-stimulation duration elapsed and the execution of the stop request. Note that completing the http POST requests introduces some temporal variability due to the execution of each step of the request, which may have slightly impacted the overall stimulation durations. However, we ascertained that the duration of requests completion, and thus stimulation durations, did not differ between active and sham modes in both conditions (all *p*s > 0.05, see Supplementary materials).

### Procedure

The study was conducted as a within-subjects, cross-over, single-blind design. Participants completed two sessions, one for the tonic and phasic conditions, scheduled at the same time of the day at least a week apart. The order of the conditions and the order of the stimulation modes within each condition were counterbalanced across participants. The second session additionally included a 5-min block of resting state pupillometry prior to the stimulated blocks, in order to assess the baseline pupil size at rest without any stimulation applied, while minimising the possibility of stimulation affecting the resting pupil size. See Fig. 1 for an illustration of the design.

In every session, participants were seated in a windowless, dimly-lit room with constant luminance. The first session began with the placement of the conductive gel or cream-coated electrode in the active position for intensity calibration. After a perceptible, but not painful, stimulation intensity level was established (see section *taVNS*), participants were seated 78 cm away from the screen, with their heads placed in a chinrest. For participants beginning the first session in the sham mode, electrode location was changed to left earlobe, and it was validated whether the established stimulation intensity is tolerable also in this location. Intensity was decreased (if painful) or increased (if not clearly perceptible) in steps of 0.2 mA. After the instructions were delivered, each block began with a 9-point eye-tracker calibration. Subsequently, participants were asked to fixate on a black-and-white fixation point, presented continuously, without offset, in the centre of a grey screen. Each block began with a 10 s baseline period (without stimulation), after which stimulation was applied in the corresponding condition/mode setting (see below). In the second session, a 5-min resting block was recorded (note that this always took place in the second session in order to equalise session durations - the resting block took comparable time as the intensity work-up procedure in session 1). Afterwards, the electrodes were placed in the first location to be stimulated, and the intensity was gradually brought to the level determined in the first session, unless rated as painful earlier or not clearly perceptible at the same level, in which case the new intensity level was maintained.

#### Phasic condition

The phasic condition followed the design of Sharon et al [36], with the stimulation duration reduced to 1 s. Eight blocks of 11 30 s-long trials were conducted, four blocks (44 trials) in the Active mode, and four blocks (44 trials) in the Sham mode (mode order was counterbalanced across participants). The first trial in each block started with a 10 s baseline period without stimulation, followed by 1 s ON-stimulation, followed by 29s OFF-stimulation. Subsequent trials in a block were separated by a jittered 800-1200ms intertrial interval.

#### Tonic condition

The tonic condition followed the classic taVNS duty cycle, 30-s on period, followed by a 30-s off period. Eight blocks of 6 60 s long trials were conducted, four blocks (24 trials) in the Active mode, and four blocks (24 trials) in the Sham mode. The first trial in each block started with a 10 s baseline period without stimulation, followed by 30 s ON-stimulation, followed by 30 s OFF-stimulation. Subsequent trials in a block were separated by a jittered 800-1200ms intertrial interval. Note that the reduced trial number compared to the phasic condition arose due to the doubled duration of each tonic trial in the 30 s on/off duty cycle.

#### Resting state measurement

The resting state measurement consisted of a single block of 11 30 s-long trials, without stimulation. Again, the first trial started with a 10 s baseline period. Subsequent trials were separated by a jittered 800-1200ms interval.

## Hypotheses

### Phasic

If phasic taVNS results in a phasic activation of the LC-NE system, we made a (pre-registered) prediction that baseline-corrected event-related pupil diameter increases following the Active ON stimulation, returning to baseline 4-8s into the off-stimulation period (following Sharon et al). Pupil diameter is predicted to be larger under Active stimulation than under Sham.

### Tonic

If tonic taVNS results in a tonic activation of the LC-NE system, we made a number of pre-registered predictions:

a. *Active v Sham, 60 s period:* Averaged baseline pupil size is larger in the entire Active condition than in Sham.
b. *Active v Sham, ON periods:* Averaged baseline pupil size is larger during Active ON period than during Sham ON period.
c. *Sham-on v resting:* There should be no significant difference in baseline pupil size in Sham ON period and resting trials. The following were formulated as research questions without directional predictions for the tonic stimulation condition:
d. *Active ON v Active OFF:* Is the effect of Active stimulation on pupil size sustained from the ON to OFF periods?
e. *Active OFF v Sham OFF:* Does the Active OFF period present a larger pupil size than Sham OFF period?
f. *Phasic-like activation, Active v Sham:* Does the onset of tonic stimulation present a phasic-like initial increase?

## Data Preparation and Analysis

### Data processing

Pupil diameter data recorded from both eyes were processed using Pupillometry Pipeliner (PuPL version 2.1.0; [47]), running on Matlab. In the first step, the portion of data, which was recorded as area, were converted to diameter following the formula: *pupil*_*diameter* = 256 * √(*pupil*_*area*/3.14159). The portion of data recorded at higher sampling frequency than 500 Hz, was downsampled to 500 Hz. All data were smoothed with a 100-ms moving mean window filter. Subsequently, blinks were detected using a velocity profile method [48]. Blinks were padded with 100 ms period pre- and post-blink, and linearly interpolated (the duration of the interpolated segment was maximum 1000 ms). Concatenated data were then segmented into epochs, depending on the condition to be analysed (phasic, tonic, and phasic-like tonic).

For the analysis of the phasic condition, data were segmented into epochs of 10 s before and 11 s after the stimulation onset. Epochs were baseline corrected by subtracting the mean of the 10-s period directly preceding each stimulation, and expressed as percent change from the baseline [49].

For the phasic-like analysis of the tonic condition, epochs of 10 s before and 30 s after the stimulation onset were extracted, baseline corrected as above, and expressed as percent change from the baseline, following the analysis of the phasic condition proper. For the further pre-registered comparisons of the tonic condition aiming to replicate previous literature [28–33], the data, to which no baseline correction or transformation was applied (not to remove any low frequency information potentially related to tonic LC activity; cf. [50]), were segmented into epochs of 30 s duration, starting from the stimulation onsets and offsets (ON periods and OFF periods respectively, see Fig. 1B).

In all analyses, epochs in which 30% or more of data were missing were excluded. Additionally, data with the largest amounts of blinks were excluded (the threshold was determined based on the visual inspection of the data). The mean number of rejected epochs in the tonic condition was 1.09 (4.54%, range: 0-17% rejected; Active: 0.51 (4.25%), Sham: 0.58 (4.83%)), and in the phasic condition 3.21 (7.30%, range 0-20% rejected; Active: 1.69 (7.68%), Sham: 1.52 (6.90%))^1^. Data is publicly available at https://osf.io/v2u9s/.

### Exclusion criteria

Participants with fewer than 50% of trials retained after blink correction and artefact rejection were flagged for removal – no such participants were present.

In the phasic condition, six subjects (out of the 58 analysable) were excluded: one could not tolerate stimulation intensity at the minimum predefined level (1 mA), and five were excluded due to either not feeling the stimulation on over 50% of blocks across the entire condition, or received a highly inconsistent stimulation intensity within any mode (e.g. cases when stimulation intensity was changed multiple times or by over 1mA during the condition, for example if participant suddenly found the stimulation too painful, or conversely, suddenly stopped feeling it), yielding a sample of 52.

In the tonic condition, three subjects were excluded: one could not tolerate stimulation intensity at the minimum predefined level (1 mA), and two were excluded due to the criteria defined above, yielding a sample of 55.

### Phasic analyses

The pre-registered analysis on the time-course of each trial was conducted with R Studio (R Core Team, 2021; RStudio Team, 2021). To compare the magnitude of pupil dilation (in percentage change over baseline) in Active versus Sham, mean pupil size values per participant per condition at each sampling point between 10 s before stimulation onset to 11 s post-stimulation were entered into paired Wilcoxon Signed Rank tests. The resulting p-values were corrected for false discovery rate (FDR) using the standard Benjamini-Hochberg (BH; [51]) method, using the *p.adjust* function from core R stats package^2^.

### Tonic analyses

For the pre-registered (and exploratory where stated) tonic comparisons, pupil sizes in arbitrary units (not baseline-corrected) were averaged into scalar values per participant per period of interest. All analyses were performed in JASP (v0.17.3; JASP Team, 2023), with default priors for the Bayes factors. For the pre-registered analysis of whether the onset of tonic stimulation causes a phasic-like initial increase in pupil size, the analysis proceeded in an identical fashion to the phasic analysis above, with mean pupil size values per participant per condition at each sampling point between 10 s before stimulation onset to 30 s post-stimulation onset (i.e. the entire ON period).

### Resting analyses

For the pre-registered (and exploratory where stated) resting measurement (no stimulation applied) comparisons to tonic condition, pupil sizes in arbitrary units (not baseline-corrected) were averaged into scalar values per participant. All analyses were performed in JASP with default priors for the Bayes factors.

## Results

### Intensity control

We successfully maintained the same intensity under Active stimulation in both the phasic and tonic conditions in 35 participants, and under Sham stimulation in 31 participants. In both conditions, the range of intensities was 1-5 mA.

During stimulation in the phasic condition, intensity in the Active mode (*M*_ACTIVE_ = 2.72 mA, *SD* = 1.33) was not significantly different from intensity in the Sham mode (*M*_SHAM_ = 2.72 mA, *SD* = 1.31; W = 32.50, *p* = 0.645, BF_10_ = 0.15). We successfully kept the intensity identical between Active and Sham modes in 41/52 subjects (in 11 participants, intensity increase or reduction was necessary once the stimulation location changed). For those participants for whom the intensity needed to be adjusted within the block (3 participants), there was also no difference between the Active and Sham modes (*M*_ΔACTIVE_ = 2.70 mA, *SD* = 1.34; *M*_ΔSHAM_ = 2.71 mA, *SD* = 1.32; W = 30.50, *p* = 0.789, BF_10_ = 0.17). 32 participants perceived Active stimulation as stronger, while 18 perceived Sham as stronger; two reported no difference.

During stimulation in the tonic condition, intensity in the Active mode (*M*_ACTIVE_ = 2.68 mA, *SD* = 1.23) was not significantly different from intensity in the Sham mode (*M*_SHAM_ = 2.64 mA, *SD* = 1.22; W = 49.00, *p* = 0.164, BF_10_ = 0.38). We successfully kept the intensity identical between Active and Sham modes in 43/55 subjects (in 12 subjects, intensity increase or reduction was necessary once the stimulation location changed). For those participants for whom the intensity needed to be adjusted within the block (3 participants), there was also no difference between the Active and Sham modes (*M*_ΔACTIVE_ = 2.67 mA, *SD* = 1.23; *M*_ΔSHAM_ = 2.64 mA, *SD* = 1.12; W = 45.50, *p* = 0.281, BF_10_ = 0.27). 32 participants perceived Active stimulation as stronger, while 21 perceived Sham as stronger; two reported no difference.

### Phasic

Pupil dilation was observed in both Active and Sham stimulation modes. Active taVNS led to a rapid pupil dilation reaching its half-maximum (2.29% over baseline) at 0.57 s after stimulation onset, peaking (4.57% over baseline) at 1.43 s, and returning back to half-maximum at 2.51 s. Pupil size returned to its average baseline value at 4.91 s. Sham taVNS provoked a similarly rapid, but smaller in magnitude, pupil dilation reaching its half-maximum (1.21% over baseline) at 0.62 s after stimulation onset, peaking (2.41% over baseline) at 1.46s, and returning back to half-maximum at 2.47 s. Pupil size returned to its average baseline value at 4.63 s. Pupil dilation was significantly greater under Active taVNS stimulation than under Sham stimulation in the time range between 0.44 s and 0.64 s, and between 0.84 s and 2.03 s post-stimulation onset (*ps* < 0.05, paired Wilcoxon signed rank tests, FDR-BH-corrected; Fig. 2A; see Supplementary Materials for an overlapping result of a cluster-based permutation test). Average pupil dilation during the significant timeframe was 3.48% (*SE* = 0.02) for Active, 1.78% (*SE* = 0.01) for Sham. There was no difference in the baseline values between Active and Sham conditions (*ps* > 0.05, paired Wilcoxon signed rank test, FDR-BH-corrected; see Fig. 2A).

**Figure 2.**
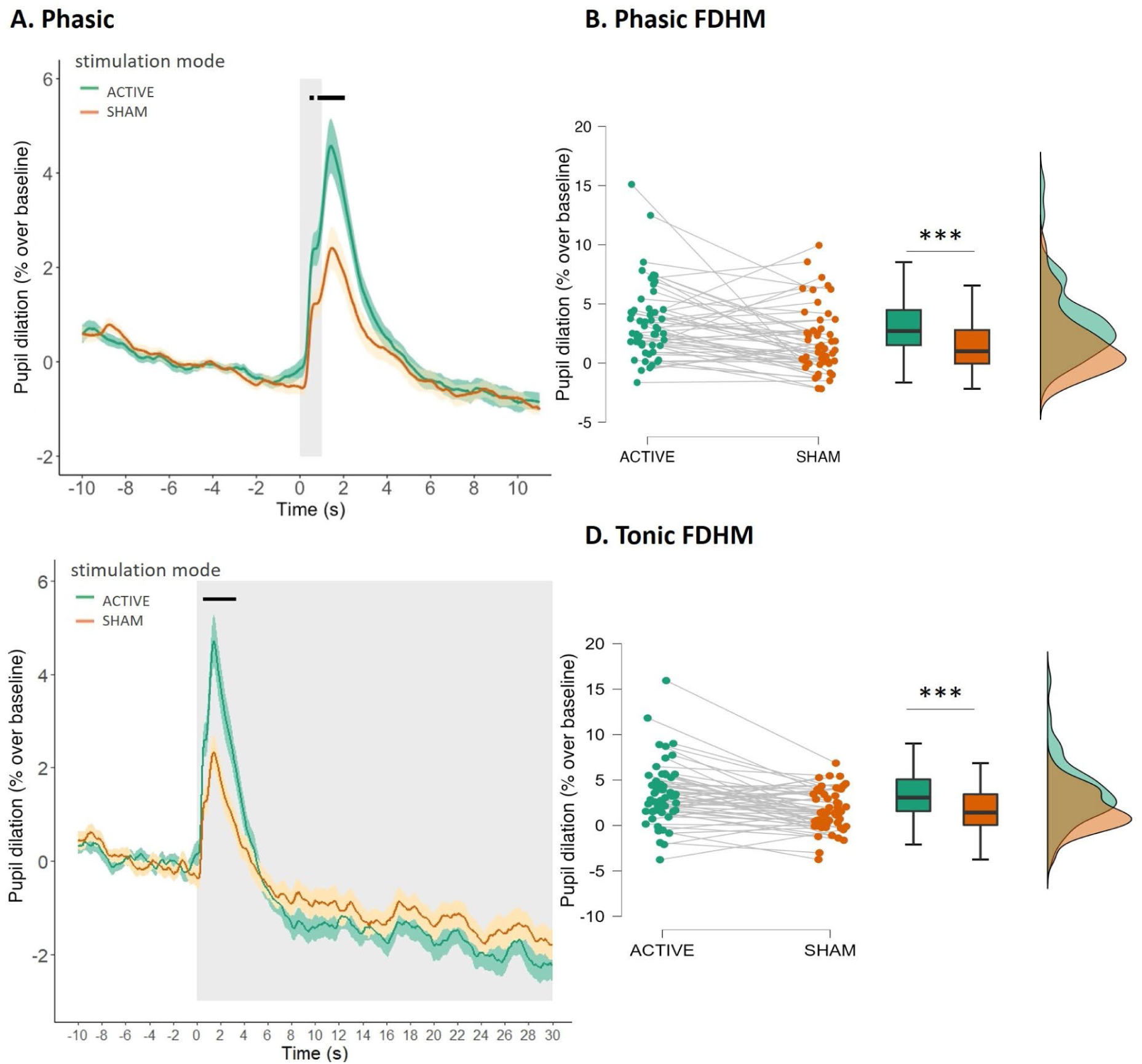
**A/C:** Pupil dilation (percentage change, baseline-corrected) in the phasic (A) and tonic (C) conditions, relative to a 10 s baseline before stimulation onset. Solid lines represent mean values. Ribbons represent SEM. The grey shading indicates stimulation duration (phasic: 1 s; tonic: 30 s, onsets without ramp-up). The short black lines indicate the period of significant difference between the values for active and sham taVNS (phasic: from 0.44 s to 0.64 s, and from 0.84 s to 2.03 s post-stimulation onset; tonic: from 0.55 s 3.36 s post-stimulation onset). **B/D:** Average pupil dilation (percentage change over baseline) for active and sham taVNS, during the full duration at half maximum, in the phasic (B) and tonic (D) conditions. In the phasic condition, 35 participants had a higher response in active stimulation mode than in sham; 17 had a higher response in sham. In the tonic condition, 42 participants had a higher response in active stimulation mode than in sham; 13 had a higher response in sham.

To further quantify the difference in pupil size between the conditions, we isolated the window of full duration at half-maximum (FDHM) from a grand average of Active and Sham time-courses (exploratory; following Sharon et al.,[36]). The pupil dilation values for active (*Mdn* = 2.70%, *SE* = 0.44, *M* = 3.44%) and sham (*Mdn* = 0.99%, *SE* = 0.39, *M* = 1.78%) were then entered into a Bayesian (default prior) paired Wilcoxon Signed Rank test in JASP. In the resulting FDHM of 0.59 – 2.50 s, active taVNS led to a significantly higher pupil dilation than sham taVNS (W = 1053, *p* < .001, BF_10_ = 45.50; Fig. 2B). Note that after removing two participants with the largest (over 10%) dilation, the difference between Active vs. Sham taVNS remained significant (W = 950.00, *p* = .003, BF_10_ = 36.30).

### Tonic: Phasic-like analysis

Pupil dilation was observed in both Active and Sham stimulation modes. Active taVNS led to rapid pupil dilation reaching its half-maximum (2.34% over baseline) at 0.55s after stimulation onset, peaking (4.67% over baseline) at 1.45s, and returning back to half-maximum at 3.04s (Fig. 2C). Pupil size returned to its average baseline value at 5.00 s. Sham taVNS provoked a similarly rapid, but smaller in magnitude, pupil dilation reaching its half-maximum (1.17% over baseline) at 0.58s after stimulation onset, peaking (2.34% over baseline) at 1.44s, and returning back to half-maximum at 2.66s. Pupil size returned to its average baseline value at 4.67 s. Pupil dilation was significantly greater under active taVNS stimulation than under sham stimulation in the time range between 0.55 – 3.36s post stimulation onset (*ps* < 0.05, paired Wilcoxon signed rank tests at each sampling point from -10 s to 30 s post-stimulation onset, FDR-BH-corrected; Fig.2C). Average pupil dilation during the significant timeframe was 3.28% (*SE* = 0.01) for active, 1.52% (*SE* = 0.01) for sham. There was no difference in the baseline values between active and sham conditions (*ps* > 0.05, FDR-BH-corrected; Fig. 2C). Early in the stimulation period, both active and sham time-courses strongly resembled the time-courses of the phasic condition, albeit peaking later, and continuing slightly longer.

To compare the average difference in pupil size between the conditions, we isolated the window of FDHM from a grand average of active and sham time-courses. The median pupil dilation values for active (*Mdn* = 3.07%, *SE* = 0.46, *M* = 3.49%) and sham (*Mdn* =1.42%, *SE* = 0.30, *M* = 1.64%) were then entered into a Bayesian (default prior) paired Wilcoxon Signed Rank test in JASP. In the resulting FDHM of 0.56 – 2.93s, active taVNS led to significantly higher pupil dilation than sham taVNS (W = 1232.00, p < .001, BF_10_ = 599.56; see Fig.2D). Note that after removing two participants with the largest (over 10%) dilation, the difference between active vs. sham taVNS remained significant (W = 1123.00, *p* < .001, BF_10_ = 212.08).

### Tonic

#### Active v Sham, 60 s period (hypothesis a.)

Averaged pupil size from the 60-s period following the onset of Active stimulation (i.e., collapsing 30 s ON and 30 s OFF periods) was compared with the pupil size in the 60-s Sham period with a paired Bayesian Wilcoxon signed-rank test. Mean pupil size values in the entire 60-s period were slightly lower for Active (*M* = 4677.83, *SE* = 75.03 a.u.) than for Sham (*M* = 4688.63, *SE* = 72.61 a.u.), and the difference was not significant (W = 732.00, *p* = 0.753; BF_10_ = 0.17; Fig. 3).

**Figure 3.**
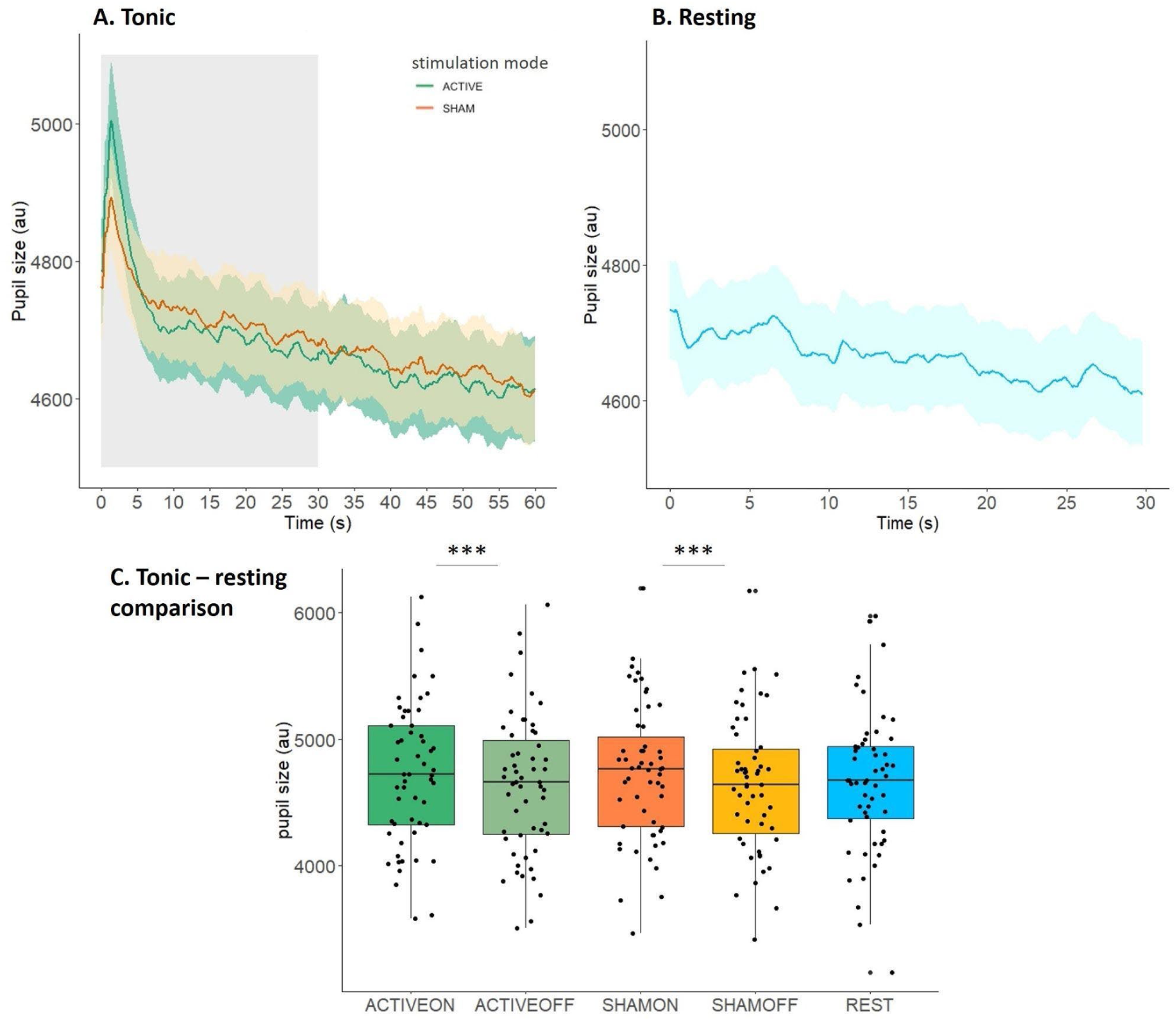
**A:** Tonic condition, averaged tonic baseline pupil sizes (a.u., no baseline correction), across the entire 60-s trial span, for Active and Sham. The grey shading indicates stimulation duration (30 s, onset without ramp-up). Ribbons represent SEM. **B:** Resting pupil measurement (no stimulation applied), averaged pupil size (non-corrected) across the 30-s blocks. Ribbons represent SEM**. C:** Comparison of pupil sizes under tonic stimulation in Active (ON/OFF) and Sham (ON/OFF) modes, and the resting measurement.

#### Active v Sham, ON periods (hyp. b.)

Averaged pupil size in the 30 s ON period of the active mode was compared to the 30 s ON period of the sham mode with a paired Bayesian Wilcoxon test. There was no difference pupil size between Active (*M* = 4724.11, *SE* = 75.68 a.u.) and Sham (*M* = 4730.60, *SE* = 73.03 a.u.; W = 736.00, *p* = 0.779; BF_10_ = 0.16).

#### Active v Sham, 8-s of the ON period (exploratory)

In order to isolate the apparent phasic-like pupil dilation, peaking and constricting rapidly after the onset of the stimulation, from potential longer-lasting and sustained changes, we compared averaged pupil sizes between Active and Sham modes in the first 8 s of the ON period with a paired Wilcoxon test (the 8-s cut-off was chosen as the point following the return to baseline). No difference in pupil size between Active (*M* = 4826.75, *SE* = 76.93 a.u.) and Sham (*M* = 4792.38, *SE* = 71.90 a.u.; W = 872.00, *p* = 0.395; BF_10_ = 0.24 was observed, in contrast to the phasic-like event-related analysis.

#### Active v Sham, 8-30s of the ON period (exploratory)

In order to isolate the potential longer - lasting shifts in pupil diameter from the transient pupil response, we then compared averaged pupil sizes between Active and Sham modes for the remainder of the ON period (8-30 s) with a paired Wilcoxon test. No difference in pupil size between Active (*M* = 4686.77, *SE* = 75.56 a.u.) and Sham (*M* = 4708.12, SE = 73.63 a.u.; W = 691.00, *p* = 0.511, BF_10_ = 0.19) were observed.

#### Active ON v Active OFF (hyp. d.)

Averaged pupil size in the 30 s Active ON period was compared with averaged pupil size in the 30 s Active OFF period, to assess whether the effect of taVNS would be sustained from the ON to OFF period. In the paired Bayesian Wilcoxon test, pupil size in Active ON (*M* = 4724.11, *SE* = 75.67 a.u.) was significantly larger than pupil size in Active OFF (*M* = 4631.55, *SE* = 74.66 a.u.; W = 1512.00, *p* < .001, BF_10_ = 59876.90), suggesting that there was a constriction of pupil size from the ON to OFF period.

#### Sham ON v Sham OFF (exploratory)

Averaged pupil size in the 30 s Sham ON period was compared with averaged pupil sizes in the 30 s Sham OFF period. In the paired Bayesian Wilcoxon test, pupil size in Sham ON (*M* = 4730.60, *SE* = 73.03 a.u.) was significantly larger than pupil size in Sham OFF (*M* = 4646.65, *SE* = 72.45 a.u.; W = 1505.00, *p* < .001, BF_10_ = 7808.77), suggesting that, similarly to Active ON vs Active OFF comparison, there was a decrease in pupil size from the ON to OFF period.

#### Active OFF v Sham OFF (hyp. e.)

Averaged pupil size in the 30 s Active OFF period was compared with averaged pupil size in the same period under Sham. In the paired Bayesian Wilcoxon test, there was no difference in pupil size values between Active OFF (*M* = 4631.55, *SE* = 74.66 a.u.) and Sham OFF (*M* = 4646.65, *SE* = 72.45 a.u.; W = 738.00, p = 0.792, BF_10_ = 0.17).

### Resting pupil size

#### Sham ON v Resting (hyp. c.)

Averaged pupil size in the 30 s Sham ON period was compared with the averaged pupil size in the 30 s resting period. In the paired Bayesian Wilcoxon test, there was no difference in pupil size between Sham ON (*M* = 4730.96, *SE* = 74.05 a.u.) and resting (*M* = 4662.85, *SE* = 74.01 a.u.; W = 585.00, *p* = 0.122, BF_10_ = 0.40), with Bayes factor providing weak support for the absence of a difference.

#### Sham ON v Resting, 0-8 s (exploratory)

Given the rapid and transient pupil dilation occurring at the beginning of tonic stimulation, averaged pupil size in the first 8 s of the Sham ON period was compared with the averaged pupil size in the first 8 s of the resting period (the 8-s cut-off was chosen as the point following the return to baseline). In the paired Bayesian Wilcoxon test, there was no difference between Sham ON (*M* = 4792.38, *SE* = 71.90 a.u.) and resting (*M* = 4703.16, *SE* = 73.47 a.u.; W = 599.00, *p* = 0.078, BF_10_ = 0.53), however, the Bayes factor does not provide sensitive evidence for the absence of a difference.

#### Active ON v Resting (exploratory)

Averaged pupil size in the 30 s Active ON period was compared with the average pupil size in the 30 s resting period. In the paired Bayesian Wilcoxon test, there was no difference in pupil size between Active ON (*M* = 4724.55, *SE* = 75.67 a.u.) and resting (*M* = 4662.85, *SE* = 74.01 a.u.; W = 602.00, *p* = 0.160, BF_10_ = 1.64), with Bayes factor providing weak support for the absence of a difference.

#### Active ON v Resting, 0-8 s (exploratory)

Similarly to the sham analysis, the averaged pupil size in the first 8 s of the Active ON period was compared with the averaged pupil size in the first 8 s of the resting period. In the paired Bayesian Wilcoxon test, the pupil size in Active ON (*M* = 4826.75, *SE* = 76.93 a.u.) was significantly larger than in the corresponding period of the resting block (*M* = 4703.16, *SE* = 73.47 a.u.; W = 495.00, *p* = 0.021, BF_10_ = 4.85), with Bayes factor providing support for the difference.

## Discussion

A better understanding of putative noradrenergic effects of taVNS is essential for refining its application and efficiency as an experimental manipulation to probe neuromodulatory influences on human behaviour. In this study, we compared the effects of phasic-like and tonic-like taVNS on pupil size, a putative biomarker associated with LC-NE activity, in a single-blind, sham-controlled within-subject design at rest – to our knowledge, the first such investigation.

In the phasic (1 s) stimulation condition, we found that active taVNS applied to left cymba conchae induced a rapid increase in pupil size over baseline while stimulation was being delivered, peaking soon after stimulation offset, and rapidly declining within two seconds. The magnitude of this increase was significantly larger than that observed after sham stimulation to the earlobe. This result corroborates recent findings that relatively brief taVNS pulses are able to elicit pupil dilation in humans [11,35–38], as well as the rodent literature showing a comparable effect of iVNS [18,39,40]. Hence, our result strengthens the claim that taVNS pulses as brief as 1 s might modulate the noradrenergic system in the phasic mode, as indexed by evoked pupil dilation.

In the tonic (30 s on/30 s off standard duty cycle) condition, matched for location and intensity, active taVNS induced a similarly rapid increase in pupil size over baseline, which was significantly larger than sham stimulation (phasic-like analysis). Importantly, however, the pupil size rapidly declined back to baseline within 5s into the ongoing stimulation, in both active and sham modes, resembling the rapid onset observed in phasic stimulation. We further unpacked the tonic effects with a pre-registered battery of comparisons on Active-Sham mean pupil size values in the ON and OFF periods. Tonic baseline pupil sizes in the whole 30-s ON period, as well as in the OFF period, did not significantly differ between Active and Sham stimulation. For both Active and Sham stimulation, pupil sizes in the ON period were significantly higher than in the OFF period, demonstrating that the increase in pupil size following stimulation was not sustained. Our analysis approach, uncovering the transient event-related effect in the tonic stimulation condition, may explain why the pre-vs post-stimulation comparisons of pupil size, or comparisons performed on binned time points, often applied to study the tonic effects of stimulation, failed to find evidence of taVNS modulation of pupil size [27,30–32]. Overall, this result suggests that prolonged taVNS under the standard set of parameters presents a phasic-like transient pupil response, with a temporal profile similar to the dilation in response to short-lived stimulation.

This result raises the question of why tonic stimulation would induce only a short-lived pupil dilation, rather than a sustained effect. An increase in pupil dilation followed by a sustained elevation over the course of the stimulation has been shown in recent animal studies using iVNS stimulation [39,40]. Such a pattern could also be expected based on an LC stimulation study with a 10-s tonic stimulation train [53], although the effect of tonic stimulation appears to manifest a parameter dependency to a larger extent than the transient pupil dilation evoked by phasic LC stimulation. Interestingly, a recent study using VNS in epileptic patients has reported that 10-s stimulation trains of different intensity levels have led to a transient pupil dilation response (albeit more sustained than control), peaking between 2.5 – 5 s, and, similar to our findings, constricting to (or even below) baseline values throughout the stimulation [54].

Similarly to previous studies [27,29–31], we also did not observe any evidence of an elevated tonic baseline - in fact, in addition to the constriction from the ON period to OFF period, we observed a steady decrease in pupil size as the block progresses, under both Active and Sham stimulation modes (see Supplementary Materials 3; note that a decrease from the beginning to the end of the task was also observed by [31], while an opposite effect was found by [32]). Similar decreases in pupil size over time have been reported previously over the course of longer trials [55], entire tasks [56], and over the course of resting measurements [28,57]. This decrease has been associated with fatigue and reduction in vigilance [58,59]. Nonetheless, it may be argued that the decline in pupil size throughout the block may have obscured potential fine-grained tonic pupil effects induced by stimulation. Although our supplementary analyses did not suggest so, this notion merits further study, as the effect may exist on a different time-scale than captured by our analyses. Finally, it is plausible that pupil dilation may be a better suited readout of phasic, rather than tonic, LC activity [9].

Given a recent report of different patterns of LC activation depending on specific combination of VNS parameters [60], another possible reason behind the absence of a sustained effect of tonic stimulation in the presence of the transient effect could be parameter choice. Firstly, it is possible that the stimulation session (24 minutes per Active/Sham mode) may have been too short to observe tonic pupil effects. Previous studies exploring tonic stimulation in humans typically use slightly longer durations (e.g. 40-55 min in D’Agostini et al. [29]). However, as described earlier, they also failed to observe the effect of prolonged stimulation on pupil size (in addition to other noradrenergic biomarkers, such as salivary alpha amylase). It could be that the effects of stimulation manifest under durations of hours or days, with compound effects of multiple stimulation sessions. Indeed, continuous 30 s on/ 5 min off iVNS in rodents delivered for 14 days showed increases in NE in the prefrontal cortex and hippocampus [61]. Elsewhere, one hour iVNS in rodents delivered at regular intervals showed a systematic increase in extracellular NE levels, which returned to baseline when not stimulated [19]. It has been proposed that such noradrenergic modulations drive the clinical effects of chronic VNS as an adjunct treatment for depression and epilepsy in humans [6–8]. Consequently, short-duration tonic taVNS might not evoke detectable shifts in the tonic LC-NE, or detectable readouts in baseline pupil size.

Secondly, it is plausible that the stimulation frequency of 25 Hz (tVNS® R default) applied in our and similar studies was well suited for a phasic effect, but ill-suited for a sustained tonic effect. In mice, increasing pulse frequencies affected the timing of LC activity, with greater increases in LC firing over a short period of time [39]. This opens up a suggestion that tonic effects on pupil size may still be found with lower frequency of stimulation over a longer period, matched to the 0.5-10 Hz frequency of tonic LC firing [20,39,62,63]. This possibility remains to be investigated. Here, we varied only the duration of stimulation (1 s vs 30 s) while keeping the other parameters constant on their standard settings (25 Hz frequency, 250 μs pulse width; notably, the intensities were matched between tonic and phasic conditions within participants as closely as possible, see next paragraph). This bolsters the notion that the interaction of stimulation duration with frequency might be an important factor in targeting the LC-NE system, with the standard setting combination potentially permitting only rapid but short-lived, phasic effects, but not sustained, tonic effects. This could be particularly important for applying phasic taVNS during behavioural tasks – although our study cannot provide a comment on the interaction between phasic stimulation and stimulus-evoked phasic responses.

Another parameter which could influence the general neuromodulatory and psychological effects of taVNS, regardless of stimulation mode, is pulse intensity. LC-NE activity readouts appear to increase monotonically with increased stimulation intensity [11,39], and there might be an intensity threshold under which LC-NE is not sufficiently activated (Helmers et al [64] propose a minimum range of 0.75-1.75 mA for iVNS, which is likely to be higher for taVNS due to skin impedance and subcutaneous tissue affecting current flow; [9]). A systematic manipulation of total charge per pulse (intensity*pulse width) revealed a dose-dependent effect of taVNS on pupil dilation, which was stronger in active than in sham stimulation, in a study using 5-s stimulation [11]. Intensity effects could also be confounded by individual differences in perceptual thresholds, and interactions with other parameters. For instance, it has been noted that perceptual thresholds may decrease with increasing pulse width [64; note tragus stimulation]. Current understanding of the interaction of objective intensity, subjective sensation, and the resulting outcomes remains limited (see [9] for a review).

It is also plausible that the phasic pupil response in both phasic-proper and at the onset of tonic stimulation does not reflect taVNS-mediated noradrenergic activity, but rather surprise or a response to a sudden intense sensation caused by the stimulation, similar to a typical stimulus-evoked phasic dilation. Indeed, somatosensory stimulation consistently evokes phasic pupil dilation [22,66, 67], which has been proposed to reflect an orienting response [68]. The orienting response itself has been proposed to rely on the LC-NE system [67,69,70], with the phasic responses observable at moderate levels of tonic discharge indicative of task responsiveness or engagement, in line with the Adaptive Gain Theory [20]. Our observation that both the phasic and tonic taVNS, in both Active and Sham modes, elicit a rapid, short-lived phasic pupil response would support the interpretation that pupil dilation could have been partly a reflection of orienting to a sudden somatosensory stimulus, regardless of stimulation location. Indeed, the similar time-course of the response to Active and Sham stimulation alike might be suggestive of the same mechanism. However, the significantly higher pupil dilation in the Active mode in comparison to Sham suggests that taVNS may still be able to induce phasic LC-NE activity over and above the orienting or somatosensory effect.

Another possibility, related to the above, is that the orienting or somatosensory effects interact with perceived intensity, visible as Active-Sham differences in stimulation-evoked pupil response. In this study, the objective intensity levels were matched as closely as possible, in an effort to keep the intensity comparable between the conditions and modes. In this vein, stimulation intensity was established for the Active mode with a subjective perception scale, and re-adjusted should participants perceive it to be either painful or poorly detectable in the Sham mode, but it was not established anew for Sham. Furthermore, for some of the participants a headband was required to secure the electrodes in the sham position. These procedural aspects may have had the unintended consequence of producing a different and/or higher perceived intensity of Active stimulation as compared to Sham, thus potentially driving a larger orienting or somatosensory effect under Active stimulation. However, in recent studies, independently established Sham intensity was typically higher than in Active mode, still leading to the same direction of the effect as found here [35,36]. In addition, dose-dependent effects of intensity were found to be stronger for Active stimulation mode than for Sham [11]. Therefore, we considered the likelihood of perceived intensity entirely driving the observed effect small. Nevertheless, to further investigate this, we analysed the interaction of perceived intensity (Active or Sham perceived as stronger) with stimulation mode (Active/Sham), finding that the main effect of mode remained significant, while a main effect of perceived intensity and its interaction with mode did not reach significance (see Supplementary material 1).

Despite taVNS driving the robust phasic pupil dilation, considered chiefly a noradrenergic biomarker, a considerable limitation to note is that taVNS is not a targeted tool - stimulation of the vagus nerve also activates other neurotransmitter systems, including the serotonergic [71], cholinergic [40], and possibly GABAergic [72] systems. However, it has been noted that the VNS-driven increase in LC firing occurs earlier than the increase in dorsal raphe (main source of serotonin) firing, which has been interpreted as serotonergic effects being secondary to the noradrenergic ones [71]. Indeed, the LC receives direct projections from the NTS, the central target of the vagus nerve [3,12,13]. Moreover, lesions to the LC abolish the noradrenergically-modulated antidepressant effects of VNS in rodents [7,73].

Despite ample evidence that LC activity is reflected in pupil size changes [21,74,75,76,77], interpreting pupil size changes as a readout of LC activity requires caution, as it is an indirect marker characterised by relatively low temporal precision [76,78]. Furthermore, one cannot isolate the noradrenergic pathway as a sole contributor to pupil diameter fluctuations. The basal forebrain-acetylcholine (BF-ACh) system has also been shown to be involved in pupil size control [23], possibly through LC projections to the BF-ACh system [76,79]. Indeed, iVNS-driven pupil dilation in rodents has been associated with cholinergic activity [40]. Noradrenergic activity was shown to be closely associated with rapid, phasic pupil dilations, while cholinergic activity was linked to longer-lasting pupil dilations at larger timescales, e.g. during locomotion [23]. Recently, transient pupil size changes have also been linked to serotonin release, particularly under conditions of low uncertainty [80]. This has been interpreted as reflecting prediction error or surprise, a signal also attributed to LC-NE [81,82], thus potentially implicating both systems in phasic (stimulus- or task-related) control of pupil size.

To conclude, in this study, we compared the effects of phasic-like (1 s) and tonic-like (30 s ON/OFF) taVNS on pupil size, a putative biomarker associated with LC-NE activity, in a single-blind, sham-controlled within-subject design at rest – to our knowledge, the first such investigation. We show that 1 s taVNS to the left cymba conchae induced a rapid and transient increase in pupil size over baseline during stimulation, which was significantly larger than under sham stimulation to the earlobe. Importantly, 30 s tonic taVNS induced a similarly rapid increase in event-related pupil size over baseline, significantly larger than under sham stimulation, resembling the transient phasic effect. No sustained effects on tonic baseline pupil size were observed. This result sheds light on the temporal profile of phasic and tonic stimulation, with implications for their applicability in further research. It suggests that the standard stimulation parameters may be better suited for phasic, rather than tonic, modulation of the noradrenergic system, and addresses the inconsistent findings in the field.

## Acknowledgements

We thank the research assistants (Alina Hansen, Ida Hoxhaj, Lea Ohanyan, Lorenzo Reuter, and Rania Lalouh) for help with data collection. We also thank Omer Sharon and Nils Kroemer for their advice, and Beth Lloyd and Franz Wurm for analysis discussion. This research was supported by the Deutsche Forschungsgemeinschaft (DFG) grant JO-787/6-1 and by the Research Foundation Flanders (FWO) postdoctoral fellowship (12V5620N).

## CRediT Author Statement

**LS:** Conceptualisation, Methodology, Software, Investigation, Data Curation, Formal Analysis, Visualisation, Writing: Original draft, Writing: Review & editing. **AM:** Conceptualisation, Methodology, Software, Investigation, Data Curation, Formal Analysis, Writing: Original draft, Writing: Review & editing. **GJ:** Conceptualisation, Funding Acquisition, Resources, Writing: Review & editing.

## Footnotes

1 Note that two pre-processing steps diverge from the pre-registered ones: choice of the filter window, and the criteria for epoch rejection. These were motivated by the availability of processing options in the PuPl tool, and are compatible in terms of reduction of high-frequency noise, and artefact rejection.

2 Note that this deviates from our pre-registered Benjamini-Yekutieli [52] method for FDR correction. Two studies published after our pre-registration [11], [35], which replicated Sharon et al findings, used a BH correction. Thus, we have opted for this correction in view of the recent publications.

