## Supplementary materials for "Tonic and phasic transcutaneous auricular vagus nerve stimulation both evoke rapid and transient pupil dilation"

#### 1. Intensity control analysis with subjective intensity differences

Despite largely keeping the stimulation intensity the same across Active and Sham modes, some participants reported perceiving Active stimulation as subjectively stronger, while others reported perceiving Sham stimulation as subjectively stronger (see *Intensity control* in the main text). Consequently, a difference in subjective sensation could affect the pupil in response to stimulation.

*Analysis*

To control for subjective sensation in both conditions (phasic, tonic), we included the reported subjective sensation in an exploratory linear mixed effects model in R (R Core Team, 2023; package *lme4*; Bates et al., 2015), with mode (Active/Sham) and subjective sensation (Active stronger / Sham stronger) as interacting predictors, and pupil dilation during the full duration at half maximum (FDHM, in percentage-change; see *Results* in the main text). Note that the two participants in each condition who reported no difference were excluded from this analysis to avoid considerable imbalance. Subject-level intercept was used. The model was fit with maximum likelihood estimation, with the following syntax: lmer(pupil size ~ mode*sensation + (1|subID), data, REML= FALSE). Analysis of deviance (Type II Wald tests) was performed on the model-estimated means using the Anova function of the *car* package (Companion to Applied Regression, Fox & Weisberg, 2019).

*Results*

*Phasic.* We found that there was a main effect of mode on pupil size during FDHM (Active/Sham; $X^2(1) = 12.33$, $p < 0.001$), corroborating the Wilcoxon result reported in the main text. We found no main effect of subjective sensation (Active stronger/Sham stronger, $X^2(1) = 1.87$, $p = 0.171$) and no interaction between mode and sensation ( $X^2(1) = 2.65$, $p = 0.103$), suggesting that subjective intensity was unlikely to modulate the observed pupil size changes (see Fig.S1).

*Tonic.* We found that there was a main effect of mode on pupil size during FDHM (Active/Sham; $X^2(1) = 20.34$, $p < 0.001$), corroborating the Wilcoxon result reported in the main text. We found no main effect of subjective sensation (Active stronger/Sham stronger, $X^2(1) = 0.14$, $p = 0.710$) and no interaction between mode and sensation ( $X^2(1) = 1.40$, $p = 0.236$), suggesting that subjective intensity was unlikely to modulate the observed pupil size changes (see Fig.S1).

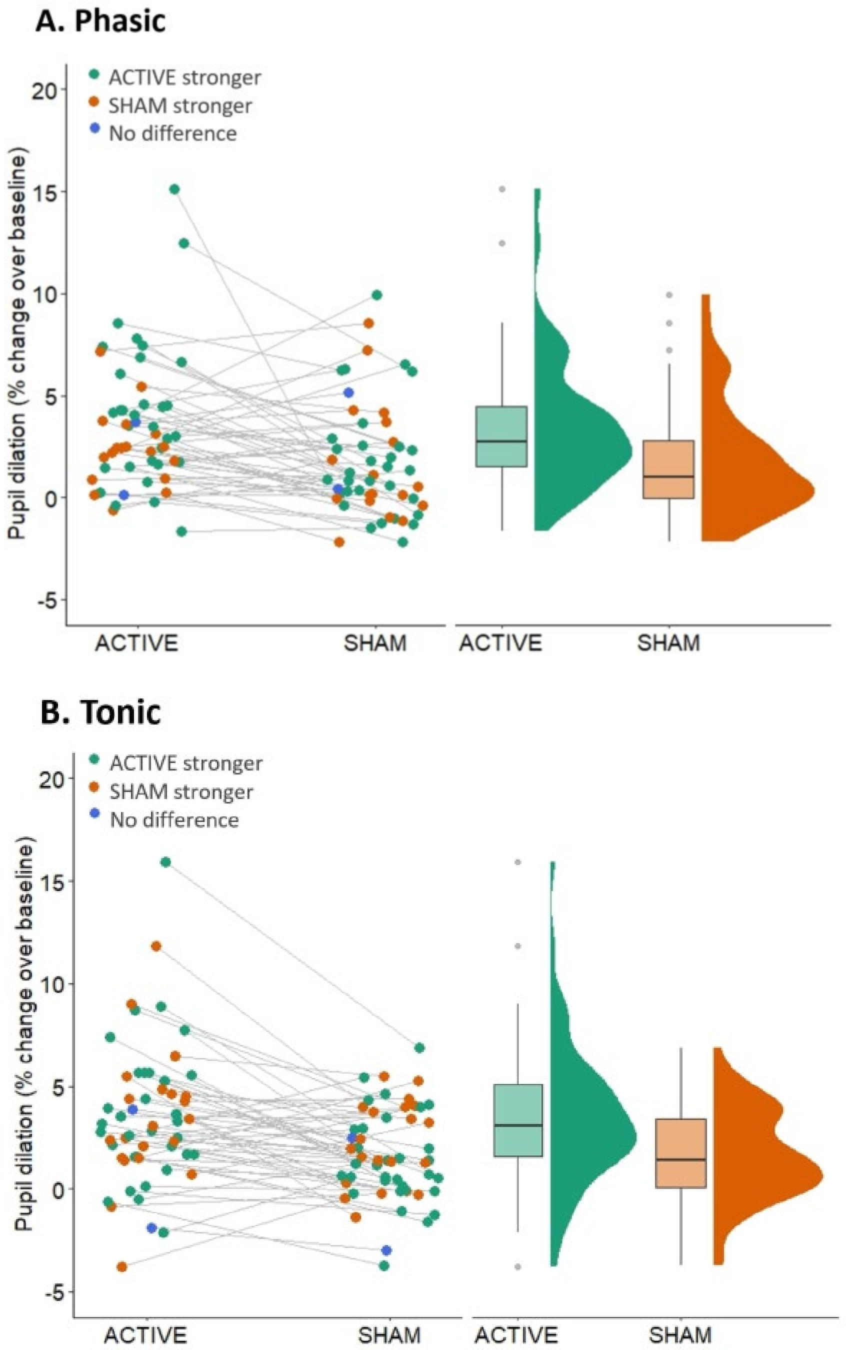


Figure S1. Pupil size in % change from baseline across full duration at half-maximum, for phasic (A.) and tonic (B.). In the dotplots, subjective sensation is presented: green dots indicate participants who reported Active stimulation as stronger, orange dots indicate participants who reported Sham stimulation as stronger, and blue dots indicate participants who reported no difference between the two. In the phasic condition, 32 participants perceived Active stimulation as stronger, while 18 perceived Sham as stronger; two reported no difference. In the tonic condition, 32 participants perceived Active stimulation as stronger, while 21 perceived Sham as stronger; two reported no difference.

### 2. Cluster-based permutation testing

In order to verify the significant periods identified through the FDR-corrected Wilcoxon tests for the phasic and tonic conditions (see main paper), we additionally performed cluster-based permutation tests (R Core Team, 2023; package *permutes*, Voeten, 2023) for the difference in pupil dilation (percentage change over baseline) between Active and Sham stimulation. For the phasic condition, the difference was significant in the periods from -8.87 to -8.27 s, from -2.56 to -2.54 s, and from -0.064 to 3.36 s. For the tonic condition, the difference was significant from -4.88 to -4.51 s; from -0.054 to 4.11 s (see Fig.S2).

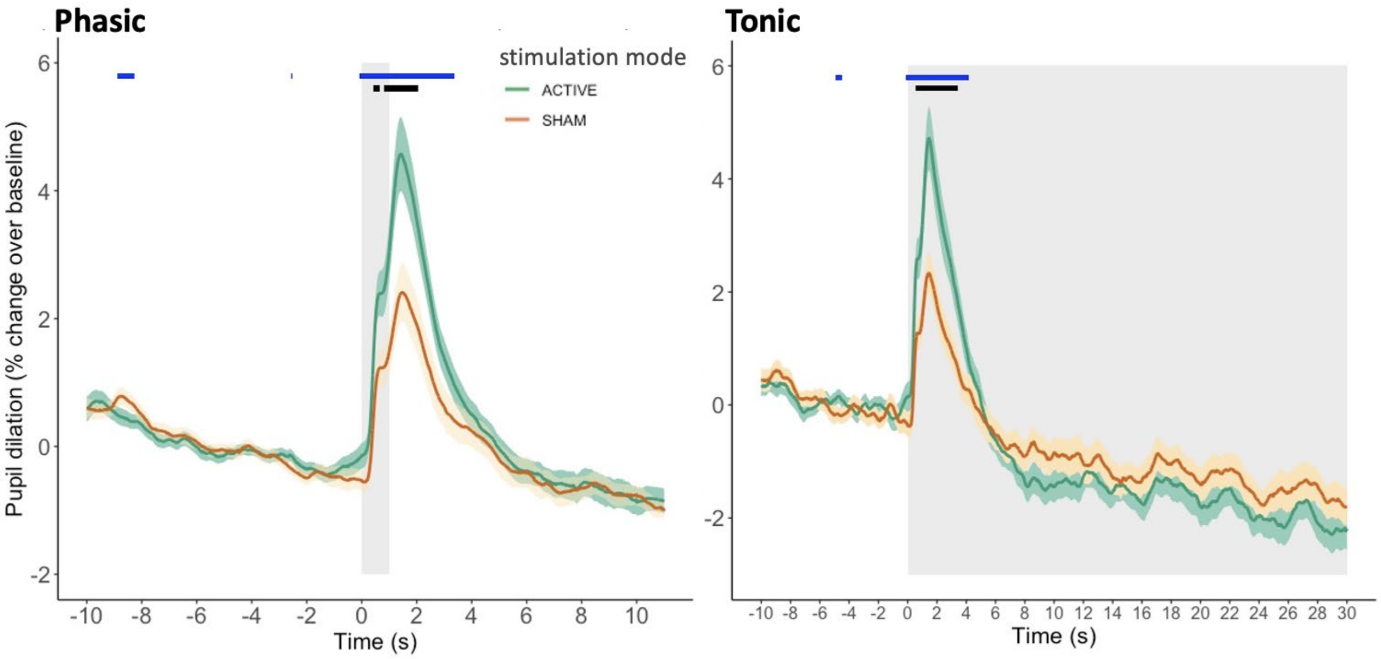


Figure S2. Pupil dilation (percentage change, baseline-corrected) in the phasic and tonic conditions, relative to a 10 s baseline before stimulation onset. Solid lines represent mean values. Ribbons represent SEM. The grey shading indicates stimulation duration (phasic: 1 s; tonic: 30 s, onsets without ramp-up). The short black lines indicate the period of significant difference between the values for active and sham taVNS (phasic: from 0.44 s to 0.64 s, and from 0.84 s to 2.03 s post-stimulation onset; tonic: from 0.55 s 3.36 s post-stimulation onset). The short blue lines indicate the identified clusters with a significant cluster mass statistic (phasic: from -8.87 to -8.27 s; from -2.56 to -2.54 s; from -0.064 to 3.36 s; tonic: from -4.88 to -4.51 s; from -0.054 to 4.11 s).

### 3. Changes in tonic baseline pupil size across the stimulation blocks.

Having observed a significant constriction, in both Active and Sham stimulation modes, from the ON-stimulation period to the OFF-stimulation period (see main paper, *Tonic* section), we concluded that the effects of stimulation on tonic baseline pupil size do not seem to be carried over to the period without stimulation. This, however, opened a question of whether the effects of stimulation may be observed on a different time-scale, such as across the trials of each block, and if they could possibly be obscured by the apparent continuous constriction of the pupil across the block. Answering this question would afford insight of greater temporal resolution into the effects of tonic stimulation on baseline pupil size.

*Analysis*

To this end, we extracted mean pupil size (not baseline-corrected) for each trial of each participant. The means were there entered as an outcome variable into an exploratory linear mixed effects model in R (R Core Team, 2023; package *lme4*; Bates et al., 2015), with mode (Active/Sham), stimulation period (ON/OFF), and trial number in block (1-6, factorised) as interacting predictors. Subject-level intercept was used. The model was fit with maximum likelihood estimation, with the following syntax: lmer(mean ~ mode*onoff*trialno + (1|subID), data, REML= FALSE). Analysis of deviance (Type II Wald tests) was performed on the model-estimated means using the Anova function of the *car* package (Companion to Applied Regression, Fox & Weisberg, 2019).

*Results*

We found that there was a main effect of stimulation period (ON/OFF; $X^2(1) = 86.14$; $p < .001$), corroborating the results of the Wilcoxon test presented in the main paper, where ON-stimulation period produced larger pupil size than the OFF-period, in both Active and Sham modes. There was no main effect of stimulation mode (Active/Sham; $X^2(1) = 1.53$, $p = 0.216$), also corroborating the reported Wilcoxon results, where raw pupil size values in the Active and Sham modes did not significantly differ.

There was a main effect of trial number ($X^2(5) = 3017.63$; $p < .001$) and a significant interaction between trial number and period ($X^2(5) = 57.77$; $p < .001$), but no significant interaction with mode ($X^2 = 1.15$, $p = 0.949$), suggesting that pupil size decreased throughout the duration of the block, in both Active and Sham conditions (see Fig.S3).

The mode-period interaction ($X^2= 0.14$, $p = 0.713$) and the three-way interaction between mode, period, and trial number ($X^2= 0.47$, $p = 0.993$) were not significant.

In addition to corroborating the results of the pre-registered tests reported in the main paper with this trial-by-trial analysis, this result shows that under both Active and Sham stimulation, pupil size decreases non-linearly as the block progresses, while the higher effect of ON-stimulation, in comparison to OFF-stimulation, is maintained until the fourth trial. This indicates that the tonic pupil baseline decreases as the block progresses, consistent with previous studies (Burger et al., 2020; Tsukahara et al., 2016; Unsworth et al., 2019). Notably, this decrease is of similar extent for both Active and Sham stimulation (cf. Burger et al., 2020).

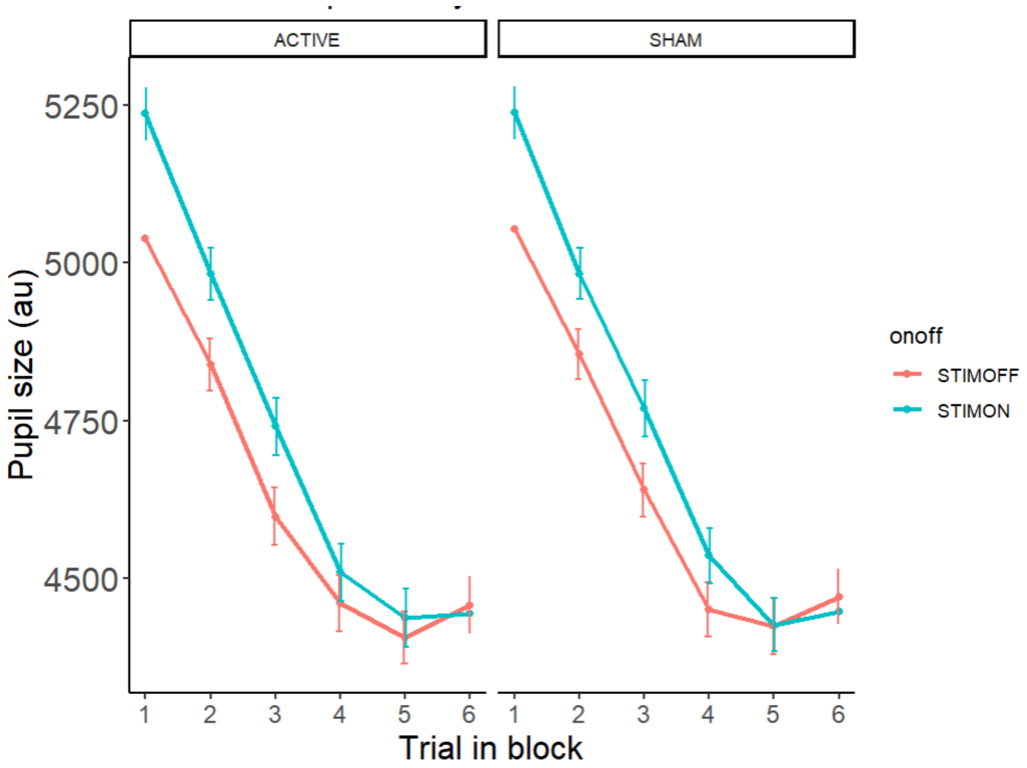


Figure S3. Pupil size (+/- SEM) difference across trials in block, for Active and Sham stimulation modes. Blue line depicts stimulation ON-periods. Red line depicts stimulation OFF-periods.

**4. Changes in baseline pupil size across the resting block.**

We subsequently assessed whether a comparable decrease in pupil size throughout the block took place during the non-stimulated resting block (conducted invariably at the beginning of session 2).

*Analysis*

We segmented the 5-min stimulation-free resting block into 10 30-s trials, comparable to the trials under tonic stimulation. We then extracted mean pupil size (not baseline-corrected) for each trial of each participant. The means were there entered as an outcome variable into an exploratory linear mixed effects model in R (R Core Team, 2023; package *lme4*; Bates et al., 2015), with trial number in the resting block (1-10, factorised). Subject-level intercept was used. The model was fit with maximum likelihood estimation, with the following syntax: lmer(mean ~ trialno + (1|subID), data,

REML= FALSE). Analysis of deviance (Type II Wald tests) was performed on the model-estimated means using the Anova function of the *car* package (Companion to Applied Regression, Fox & Weisberg, 2011).

*Results*

There was a main effect of trial number in the resting block ($X^2(9)$ = 524.51, $p$ < 0.001), suggesting that pupil size significantly decreased throughout the resting block (see Fig.S4), similarly to stimulation blocks.

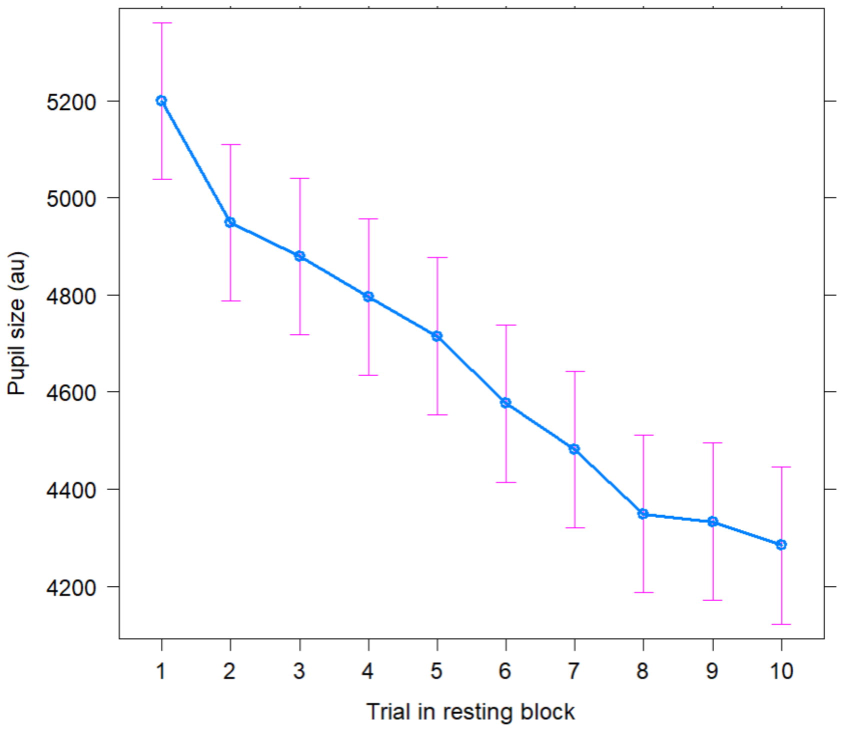


Figure S4. Pupil size (95% CI) across trials in the stimulation-free resting block.

### 5. Function duration comparisons

In order to ascertain that the durations of HTTP POST request to the tVNS Manager app (see section *Stimuli and Equipment, taVNS)* did not differ between the active and sham stimulation modes, we compared the respective durations with paired-sample t-tests, in both conditions (phasic, tonic). The durations of execution of the complete request did not significantly differ between active and sham, for both the ON- and OFF-commands (see Table S1).

**Table S1.** Comparison of durations of execution of the ON/OFF https POST requests used to control the onsets and offsets of stimulation.

| | | Active *M(SE)* | Sham *M(SE)* | Comparison |
|---|---|---|---|---|
| **Phasic** | ON-command | 1.47(0.04) s | 1.41(0.03) s | t(51) = 1.17, $p$ = 0.249, $BF_{10}$ = 0.29 |
| | OFF-command | 0.27(0.01) s | 0.27(0.01) s | t(51) = -0.15, $p$ = 0.886, $BF_{10}$ = 0.15 |
| **Tonic** | ON-command | 1.41(0.06) s | 1.35(0.05) s | t(54) = 0.95, $p$ = 0.349, $BF_{10}$ = 0.23 |
| | OFF-command | 1.59(0.06) s | 1.49(0.05) s | t(54) = 1.38, $p$ = 0.173, $BF_{10}$ = 0.36 |
